# Maternal age alters offspring lifespan, fitness, and lifespan extension under caloric restriction

**DOI:** 10.1101/527770

**Authors:** Martha J. Bock, George C. Jarvis, Emily L. Corey, Emily E. Stone, Kristin E. Gribble

## Abstract

Maternal age has a negative effect on offspring lifespan in a range of taxa and is hypothesized to influence the evolution of aging. However, the mechanisms of maternal age effects are unknown, and it remains unclear if maternal age alters offspring response to therapeutic interventions to aging. Here, we evaluate maternal age effects on offspring lifespan, reproduction, and the response to caloric restriction, and investigate maternal investment as a source of maternal age effects using the rotifer, *Brachionus manjavacas*, an aquatic invertebrate. We found that offspring lifespan and fecundity decline with increasing maternal age. Caloric restriction increases lifespan in all offspring, but the magnitude of lifespan extension is greater in the offspring from older mothers. The trade-off between reproduction and lifespan extension under low food conditions expected by life history theory is observed in young-mother offspring, but not in old-mother offspring. Age-related changes in maternal resource allocation to reproduction do not drive changes in offspring fitness or plasticity under caloric restriction in *B. manjavacas*. Our results suggest that the declines in reproduction in old-mother offspring negate the evolutionary fitness benefits of lifespan extension under caloric restriction.

## INTRODUCTION

Maternal effects occur when the environment or physiological state of a mother changes the phenotype of her offspring without a corresponding change in genotype. Offspring phenotype may be modified in response to maternal environmental factors including diet, temperature, or exposure to stressors ^1-11^. Such maternal effects may be adaptive as in *Daphnia* and rotifers, in which offspring hatch with protective spines upon maternal exposure to predators^2-6,7^, or as in plants, in which offspring have higher rates of germination and survival when planted in the same high-light or low-light environment as their parent ^7,8^. Alternatively, maternal effects may be detrimental as is the case in the negative health outcomes for children due to excessive maternal smoking or alcohol consumption during pregnancy ^13-15^. We are beginning to understand that maternal effects may be mediated by a variety of epigenetic mechanisms, including direct transmission of maternal proteins, mRNA, lncRNA, miRNA, and modifications to DNA and histones ^16-20^. While maternal effects have long been studied and are well known in the ecological literature, there has been a recent rise in interest in maternal effects in the context of human health and aging ^21^.

Maternal age, or the age of a mother at the time her offspring are born, has been shown to have a negative effect on offspring health in a range of taxa ^22-32^. A decrease in offspring lifespan with increasing maternal age was first demonstrated in rotifers--microscopic, aquatic invertebrate animals--and has come to be known as the “Lansing Effect” ^23,24,33^. Declines in offspring health, lifespan, and stress resistance with increasing maternal age have since been demonstrated across taxa, ranging from invertebrates like soil mites and *Drosophila*, to mammals including mice and humans ^26,27,29-32,34-36^. The mechanisms of these maternal age effects are unclear, and have variously been attributed to increases or decreases in maternal investment in reproduction with increasing maternal age, as well as to other, as yet undefined, epigenetic factors ^37-44^.

Epidemiological and demographic studies in humans have shown a negative correlation between maternal age and children’s lifespan and health ^25,27,30,45,46^. However, maternal age effects in humans can be difficult to separate from confounding environmental factors including paternal age effects, parental health, parental socio-economic status, and parental care ^47-49^. Additionally, in human studies, both genotype and environment are usually uncharacterized, and it is impossible to systematically and simultaneously vary maternal age and offspring environment for a given genotype. Given these challenges, appropriate animal models must be used to characterize the drivers and outcomes of maternal age effects on offspring fitness in varied environments.

Maternal effects result in different outcomes in diverse offspring environments. For example, maternal effects may be detrimental as in the Barker Hypothesis, where fetal undernutrition reprograms offspring to have a more efficient metabolism. This maternal effect is adaptive in low nutrient environments (the “thrifty phenotype”), but becomes maladaptive when children mature in high food environments, leading to adult metabolic and cardiac disease ^9-11^. Thus, maternal age may modulate the effectiveness of anti-aging lifestyle or medical interventions in offspring in unforeseen ways. While there has been some investigation of how genetic background may affect the response to lifespan-extending interventions such as caloric restriction, gene knockdown, or pharmaceuticals ^50-53^, little is known about how maternal age may influence offspring response to these therapies, or if such interventions might rescue offspring from the negative effects of maternal age.

Caloric restriction—a decrease in food consumption—has been shown to extend lifespan across a range of taxa and is heavily studied as a therapeutic intervention to aging ^50,54-58^. Evolutionary life history theory and the related Disposable Soma theory of aging both hypothesize that under the low food conditions of caloric restriction, an individual re-allocates resources from reproduction and dedicates them to preservation of the body, or soma^55,59-62^. Although it is known that maternal age affects offspring phenotype, current evolutionary theories of aging and caloric restriction do not incorporate maternal age as a variable, and thus do not describe or predict changes in the direction or magnitude of lifespan and reproductive trade-offs due to maternal age ^60,62-65^. Given the emphasis on caloric restriction and caloric restriction mimetics as interventions to increase lifespan and improve late-age health, it is critical to understand sources of variability such as maternal age in the lifespan and health responses to these therapies.

The influence of maternal age on offspring evolutionary fitness and on the evolution of aging remains poorly understood ^29,66-68^. Offspring lifespan is often measured in studies of maternal age effects, but is only one component of evolutionary fitness. To understand what drives the evolution of aging and the response to therapies, we must consider the combination of factors that contribute to fitness, including lifespan, reproduction, and resistance to external mortality as age-specific rather than as end-point traits like median lifespan and lifetime reproduction trade-offs.

In this study, we used the monogonont rotifer, *Brachionus manjavacas,* to investigate the effect of maternal age on offspring lifespan and fitness under fully fed and anti-aging caloric restriction diets. With a short lifespan of two weeks and simple laboratory culture, rotifers are similar to other tractable invertebrate model systems relevant to human health ^69,70^. In addition, rotifers provide a number of unique benefits as a model system for aging and maternal effects. *Brachionus manjavacas*, like humans, makes a relatively large investment in individual offspring, as evidenced by the low numbers of offspring produced over the two-week lifespan (25 – 30 offspring) and large egg size (30 – 50 % of adult body size). In contrast, other invertebrates, such as *C. elegans* and *Drosophila,* produce hundreds to thousands of small eggs per individual. Additionally, reproduction in *B. manjavacas* is continuous and sequential throughout the reproductive period, unlike in *C. elegans, Drosophila,* or *Daphnia,* which produce hundreds of eggs over just a few days or in clutches ^71-74^. In these ways, the reproductive strategy of *B. manjavacas* is akin to that of K-selected species like humans, rather than to r-selected species like *C. elegans, Drosophila,* or *Daphnia* ^75^. Such differences in reproductive strategies are likely to influence maternal effects on offspring. Monogonont rotifers exhibit no post-hatching parental care, avoiding the confounding effects of changes in maternal care with increasing age. Similar to humans, rotifers have direct development, with no larval stage or metamorphosis.

To eliminate confounding variability introduced by paternal effects, mother-offspring conflict, genetic recombination, and genotype diversity, we used a clonal, asexual female lineage of *B. manjavacas*. *Brachionus* spp. generally reproduce asexually, with females producing isogenic offspring via mitosis in the germline. In response to environmental conditions like crowding, some females become sexual and produce haploid male offspring that mate with other sexual females. Asexual females and their offspring were used in all experiments except for some measures of maternal investment, for which we examined meiotically-produced eggs that hatch into males. All offspring were from the same group of mothers, not from different cohorts for each maternal age as in many other studies of maternal effects; the F1 maternal age and diet cohorts were genetically-identical and composed of sets of siblings ^29^. All observations were made on individuals, not populations or groups of rotifers, and thus we can directly correlate lifespans and fecundities of individual mothers and their daughters, allowing examination of possible individual heritability of lifespan.

Because maternal environment and physiology are known to affect offspring phenotype, and because maternal age is known to influence offspring lifespan, we hypothesized that maternal age may affect offspring adaptive response to caloric restriction. This study expands upon our prior work demonstrating that maternal caloric restriction increases offspring lifespan and reproduction, especially in late maternal age offspring ^76^. In the current study, we investigated the combined effect of maternal age and offspring diet to determine (1) whether changes in gross maternal reproductive investment with increasing maternal age are correlated with offspring survivorship; (2) the extent to which increasing maternal age changes offspring response to the well-studied anti-aging therapy of caloric restriction; and (3) how maternal age and offspring diet interact to determine offspring relative age-specific reproduction as a measure of evolutionary fitness.

## MATERIALS AND METHODS

### Rotifer and phytoplankton culture

We used the Russian strain of the monogonont rotifer *Brachionus manjavacas* (BmanRUS) in all experiments. Rotifers were fed the chlorophyte algae *Tetraselmis suecica*, which was maintained in semi-continuous culture in bubbled 2-L flasks of f/2 medium^77^, made with 15ppt Instant Ocean (Instant Ocean Spectrum Brands, Blacksburg, VA). We cultured rotifers and algae at 21 °C under cool-white fluorescent bulbs at an intensity of 100 μE m^-2^s^-1^ on a 12:12 h light:dark cycle.

### Offspring lifespan, fecundity, and response to caloric restriction

We conducted life table experiments as previously described ^78^. To avoid residual undefined parental effects on our experimental populations, we synchronized the maternal ages of the great-grand and grand-maternal generations for the experimental maternal (F0) cohort by collecting eggs from 3 – 5 d old females for two generations. Briefly, we harvested eggs from a batch culture by vortexing and micropipette isolation, let these hatch and then grow in *ad libitum T. suecica* (AL; 6 × 10^5^ cells ml^-1^) for 5 days, and then collected the eggs from that culture. This was repeated twice, so that the maternal and grand-maternal ages for our experimental F0 cohort were 3 – 5 days old.

To obtain the F0 generation, eggs from this age-synchronized culture were harvested as above, allowed to hatch over 16 hours, and neonates were randomly deposited individually into 1 ml of 15 ppt seawater and AL *T. suecica* in wells of unshaken 24-well tissue culture plates (n=187). Every 24 h, we recorded survival, reproductive status (whether carrying eggs), and the number of live offspring and unhatched, dropped eggs for each individual; the female was then transferred to a new well with fresh algae of the appropriate concentration. To obtain the F1 cohorts, at the specified maternal ages we isolated one female neonate hatched within the previous 24 h from each F0 female and placed these in wells of unshaken 24-well plates with 1 ml of the appropriate food concentration (n = 69 - 72 for each F0 age X F1 diet cohort). All F1 cohorts were collected from the same set of 187 mothers. Offspring were randomly distributed among food treatments. We tested for effects of non-independence of offspring lifespan using linear regression and found no correlation in lifespan between individual F0s and their offspring for any maternal age or diet cohort. Sample size was determined by power analysis to detect a 0.75 d (approx. 7%) difference in lifespan using the program G*Power ^79^.

As a measure of offspring ability to mount a beneficial adaptive response, we subjected F1 individuals from 3, 5, 7, and 9 d old mothers (F1_3_, F1_5_, F1_7_, and F1_9_, respectively) to either chronic caloric restriction (CCR; 10% of AL food levels; 6 × 10^4^ cells ml^-1^) or intermittent fasting (IF; feeding AL or starving every other day), two treatments known to increase lifespan in rotifers (experimental design in Fig. 1). Survival, reproductive status, and numbers of offspring and unhatched eggs were recorded every 24 hours for the caloric restriction experiments. No blinding was used.

**Fig 1.**
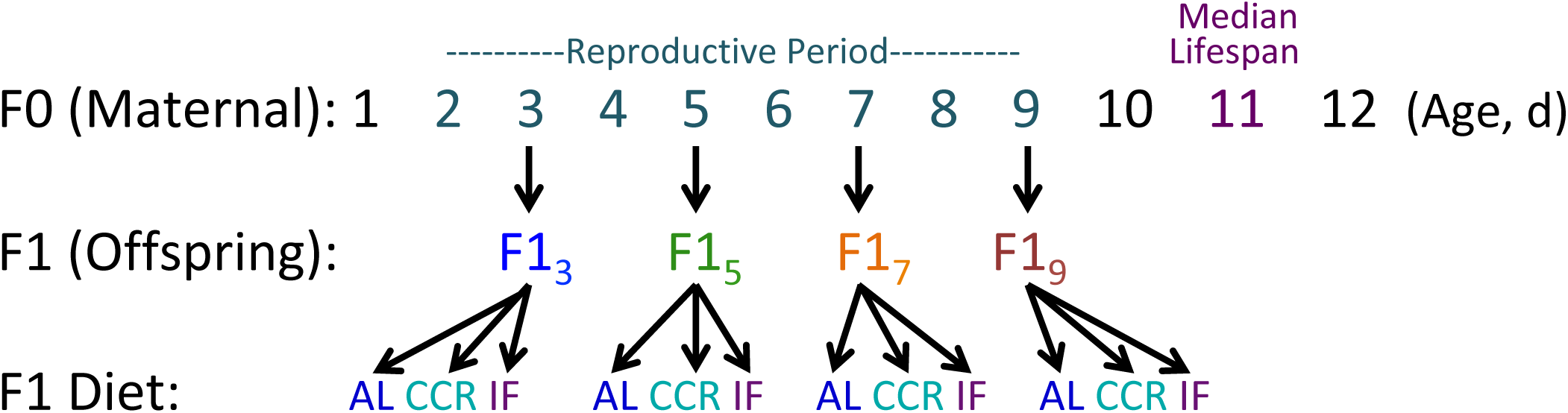
Experimental design to test the combined effects of maternal age and offspring diet on offspring lifespan and fecundity. Newly-hatched offspring (F1) were collected from *ad libitum* fed, age-synchronized amictic (asexual) maternal females (F0, n = 180) at maternal ages of 3, 5, 7, and 9 days (F1_3_, F1_5_, F1_7_, and F1_9_, respectively). Offspring were subjected to an *ad libitum* diet (AL; 6 × 10^5^ cells ml^-1^ *Tetraselmis suecica*), chronic caloric restriction (CCR; 6 × 10^4^ cells ml^-1^ *T. suecica*, a 90% reduction in food relative to AL); or intermittent fasting (IF; alternate day AL and starvation). All rotifers were housed individually in 1 ml 15 ppt Instant Ocean and algae in 24-well plates. Survival and reproduction of the F0 and F1 were scored daily until all rotifers had died. For each F1 maternal age X diet cohort, n = 69 - 72.

### Maternal investment

To determine if maternal investment in reproduction changes with maternal age, we conducted a separate experiment to measure size and shape of female and male eggs from 3, 6, 9, and 11 d old mothers. Age-synchronized females were placed 2 per well in 1 ml of 6 × 10^5^ cells ml^-1^ *T. suecica* in 15 ppt Instant Ocean in 24-well plates, and transferred daily to new wells with fresh *T. suecica.* At the specified ages, 48 – 72 rotifers were collected and vortexed or sheared through a 23 gauge needle to separate eggs (normally carried externally by females until hatching) from females. For egg size and shape, we fixed samples in 5% formalin (final concentration). Before imaging, formalin was removed by centrifugation and aspiration, and eggs were washed twice with Instant Ocean. At least 25 each of male and female eggs were imaged with a Zeiss AxioCam at 400X magnification on an Axioskop (Carl Zeiss, Inc., Thornwood, NY). We measured egg diameter, area, and roundness (inverse of aspect ratio between longest and shortest axes) using the image analysis software, Fiji ^80^.

As a quantitative assessment of changes in nutrient allocation to offspring, in a separate experiment we measured neutral lipids in newly hatched F1 neonates from 3, 6, 9, and 11 d old mothers. These lipids are maternally distributed to offspring and used as a source of nutrition by neonates post-hatching. We anesthetized 6-h old neonates in 1.0 μM bupivicain for 10 minutes before fixation in 2.5% formalin. Neonates were stained with 0.5 μg μl^-1^ Nile Red in acetone for 5 minutes and washed twice with 15 ppt Instant Ocean. For each maternal age, we imaged 20 stained and 5 unstained neonates at 200X with a Zeiss LSM 710 Confocal Microscope (Carl Zeiss, Inc., Thornwood, NY) using a 514 nm laser excitation with 559-621 nm emission and a 458/514 nm main beam splitter, imaging the entire animal volume with 1 μM slices. Lipid volume per animal volume was quantified using Fiji ^80^.

For an additional estimate of changes in maternal investment in reproduction with increasing maternal age, we measured resistance to starvation in unfed F1s from 3, 6, 9 and 11 d old mothers in a separate experiment. We isolated eggs from mothers as described above. Eggs hatched overnight in 15 ppt Instant Ocean, so that neonates were never fed, after which we placed 2 neonates per well in 1 ml of 15 ppt Instant Ocean in 24-well plates (n = 48 for each maternal age cohort). As above, sample size was determined by power analysis to detect a difference in lifespan of 0.75 d between groups ^79^. We scored survival twice per 24 hours, at 8 and 16-hour intervals, until all individuals had died.

### Statistical analyses

We used Prism 7.0a for graphing and statistical analyses. From lifespan data, we calculated median and maximum (age of 5% survivorship) lifespan. Kaplan Meier survivorship curves were constructed from lifespan data; data were right-censored in the event an individual was lost prior to death or due to accidental death caused by mishandling. Significance of differences between median lifespans was calculated using a Mantel-Cox log-rank test. We used ANCOVA to determine significance of differences between mortality rate (the slope, β) and onset of senescence (the intercept, α) from a Gompertz function fitted to age-specific hazard rate. We used one-way ANOVA with Tukey’s test for multiple comparisons to determine significant differences between egg size, shape, and lipid content across maternal ages. We used two-way ANOVA with Tukey’s test for multiple comparisons to determine significant differences in lifetime reproduction, non-viable embryos, or reproductive period between F1s due to maternal age or F1 diet. To determine the effect of interaction between maternal age and F1 diet on F1 lifespan and F1 reproduction, we used two-way ANOVA. Correlations between lifespan and reproduction were fit with a second-order polynomial (quadratic) equation; we also tested linear and third-order polymomial regressions, and found quadratic equations to be the best fit. Differences between reproduction-lifespan correlations were determined using an extra-sum-of-squares F-test.

## RESULTS

### Offspring Lifespan

Both maternal age and offspring diet affected offspring lifespan (Fig. 2, Fig. 3). Median offspring lifespan declined significantly with increasing maternal age, but maximum lifespan did not change (Supplementary Table 1). Both CCR and IF significantly increased median and maximum lifespan in all F1s, regardless of maternal age (Fig. 3, Supplementary Table 1). Under CCR, the percent increase in median lifespan was greater for F1_5_ - F1_9_ than for F1_3_, and under IF it was greater for F1_7_ and F1_9_ than for F1_3_ and F1_5_ (Supplementary Table 1). There was no significant correlation between lifespans of individual mothers and their offspring for any maternal age or under any F1 diet (data not shown, R^2^ < 0.07 for all linear regressions). There was a significant interaction between maternal age and offspring diet to determine F1 lifespan (4.26% of total variance, F_6, 791_ = 7.33, p < 0.0001). F1 diet had a greater influence on F1 lifespan (10.61% of total variance, F_2, 791_ = 54.78, p < 0.0001) than did F0 age (F_3, 791_ = 26.66, p < 0.0001).

**Fig 2.**
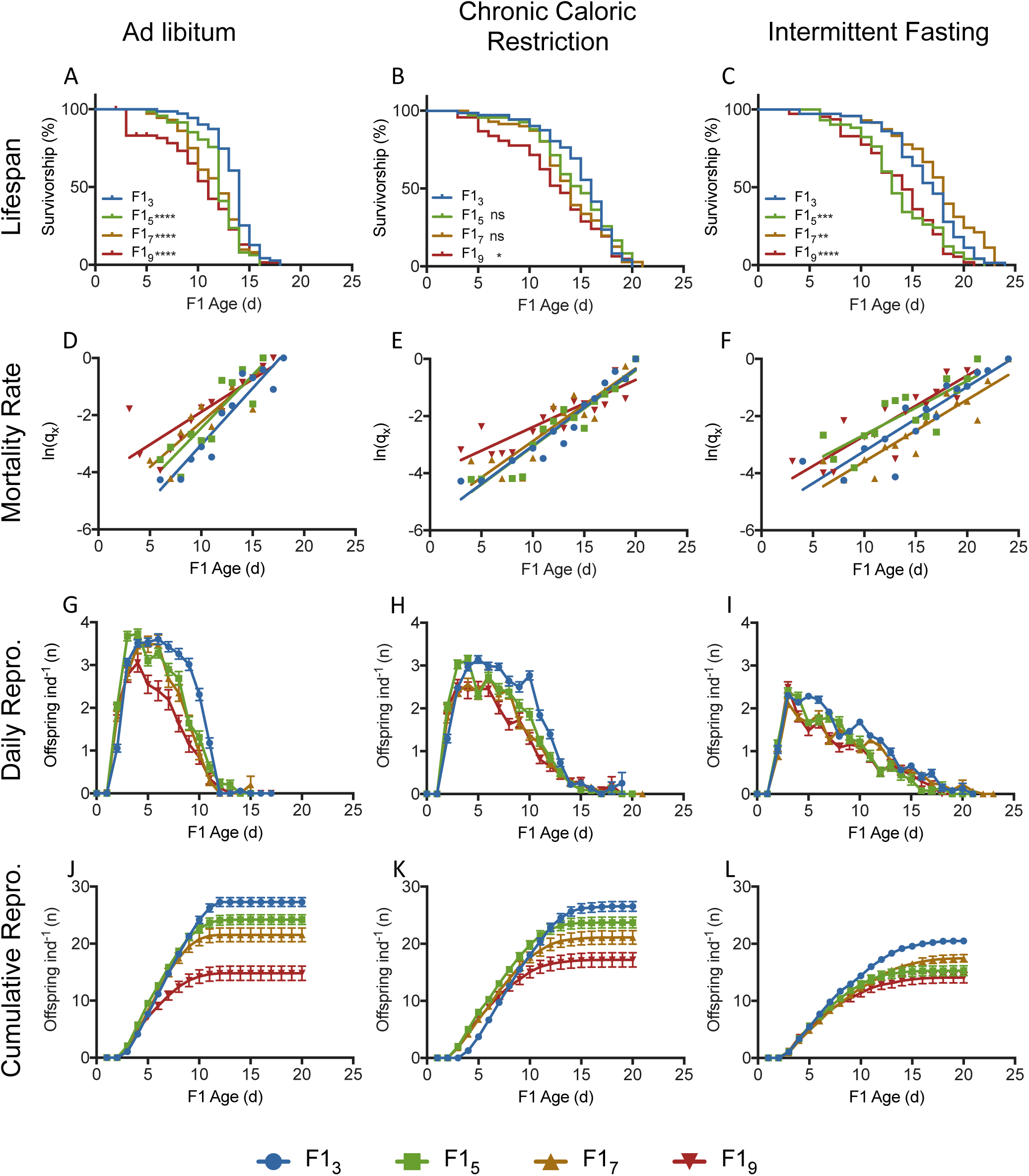
Lifespan and fecundity of F1s from 3, 5, 7 and 9 d old mothers (F1_3_, F1_5_, F1_7_, and F1_9_, respectively) under different diets. Survivorship (**A-C**), hazard rate (**D-F**), daily fecundity (**G-I**), and cumulative fecundity (**J-L**) for F1s fed under *ad libitum* (AL; **A, D, G, J**), chronic caloric restriction (CCR; **B, E, H, K**) or intermittent fasting (IF; **C, F, I, L**) conditions. Significant differences from F1_3_ are given by * (p < 0.05), ** (p < 0.01), *** (p < 0.001), or **** (p < 0.0001), or ns = not significant. Additional significance of differences in survivorship and hazard rate is given in Supplementary Table 1. Statistical significance of differences in reproduction is shown in Supplementary Table 2. For each F1 maternal age X diet cohort, n = 69 – 72.

**Fig 3.**
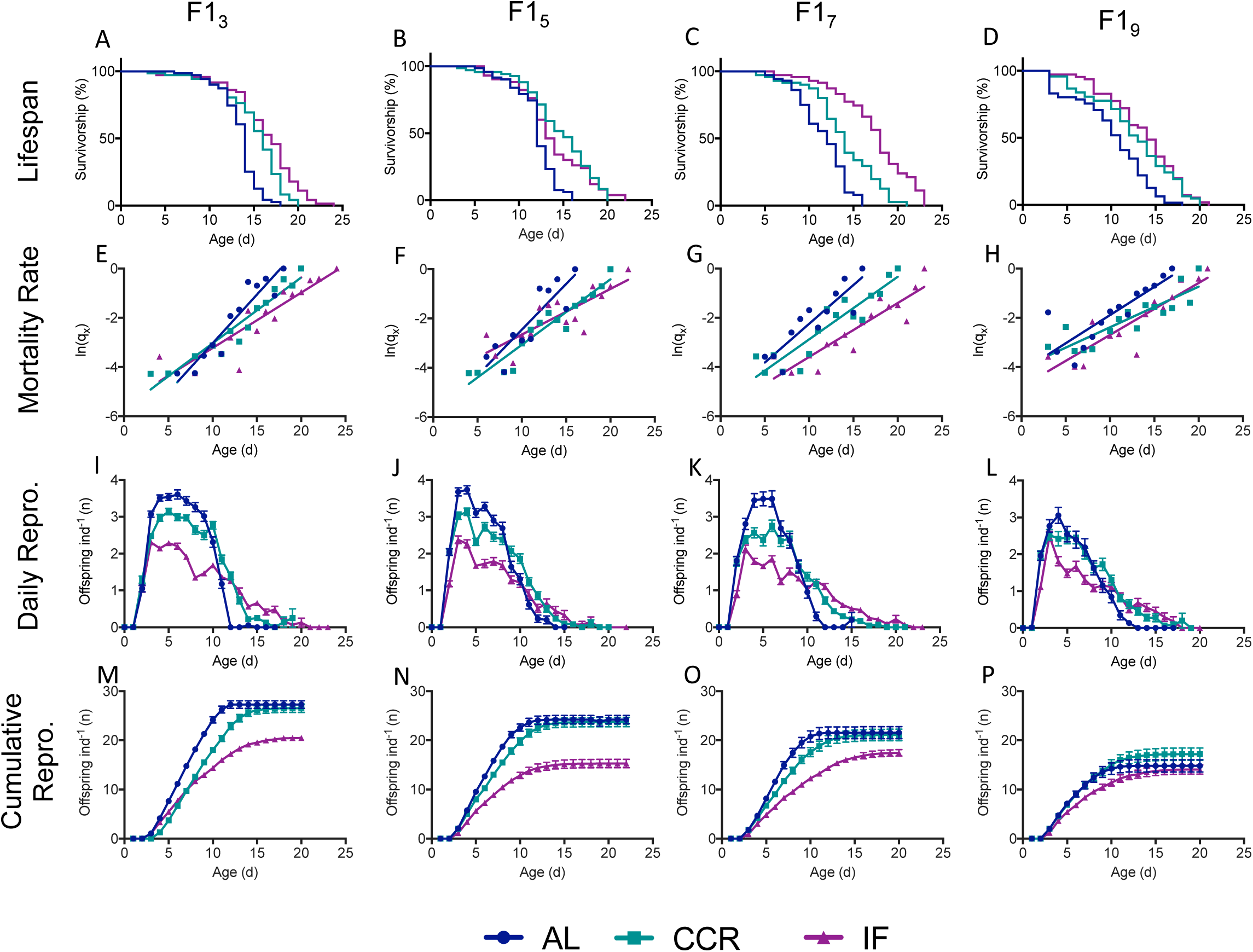
Lifespan and fecundity of F1s under different caloric restriction diets from mothers of different ages. Survivorship (**A-D**), hazard rate (**E-H**), daily fecundity (**I-L**), and cumulative fecundity (**M-P**) for F1s from 3, 5, 7, or 9 d old mothers (F1_3_, F1_5_, F1_7_, and F1_9_, respectively). n = 69 - 72 for each F1 maternal age X diet cohort. Statistical significance of differences in survivorship and hazard rate is given in Supplementary Table 1. Statistical significance of differences in reproduction is shown in Supplementary Table 2.

We estimated the onset and rate of aging by fitting a Gompertz function to the age-specific hazard rate (Fig. 2 D-F). Linear regression showed a change in both onset and rate of mortality under caloric restriction (Fig. 3), though this varied depending on maternal age (Fig. 2; Supplementary Table 1). The onset of mortality (α) was delayed under CCR and IF for offspring of all maternal ages. For younger maternal ages (F1_3_ and F1_5_), the onset of mortality was later under IF than CCR, while the reverse was true for offspring from later maternal ages (F1_7_ and F1_9_). While the rate of aging (β) was significantly lower under both CCR and IF for F1_3_, at later maternal ages there was no significant difference between the Gompertz regression slopes under AL and CCR or IF, and differences in lifespan under caloric restriction were primarily due to decreased mortality at early ages, rather than a decline in the rate of aging.

### Offspring fecundity and reproductive schedule

Increasing maternal age suppressed daily and total reproduction in the F1 (Figs. 2, 3, 4, Supplementary Table 2). While F1 CCR slightly depressed daily reproduction relative to AL, the reproductive period was extended, resulting in the same lifetime reproduction (Supplementary Table 3, Figs. 3, 4). Although the reproductive period was also extended under IF (Supplementary Table 3, Fig. 4), daily reproduction was approximately half that of AL, and discontinuous reproduction (days of no reproduction interrupting the reproductive period) was significantly higher (Supplementary Table 4), leading to significantly lower lifetime reproduction under IF (Fig 4).

**Fig 4.**
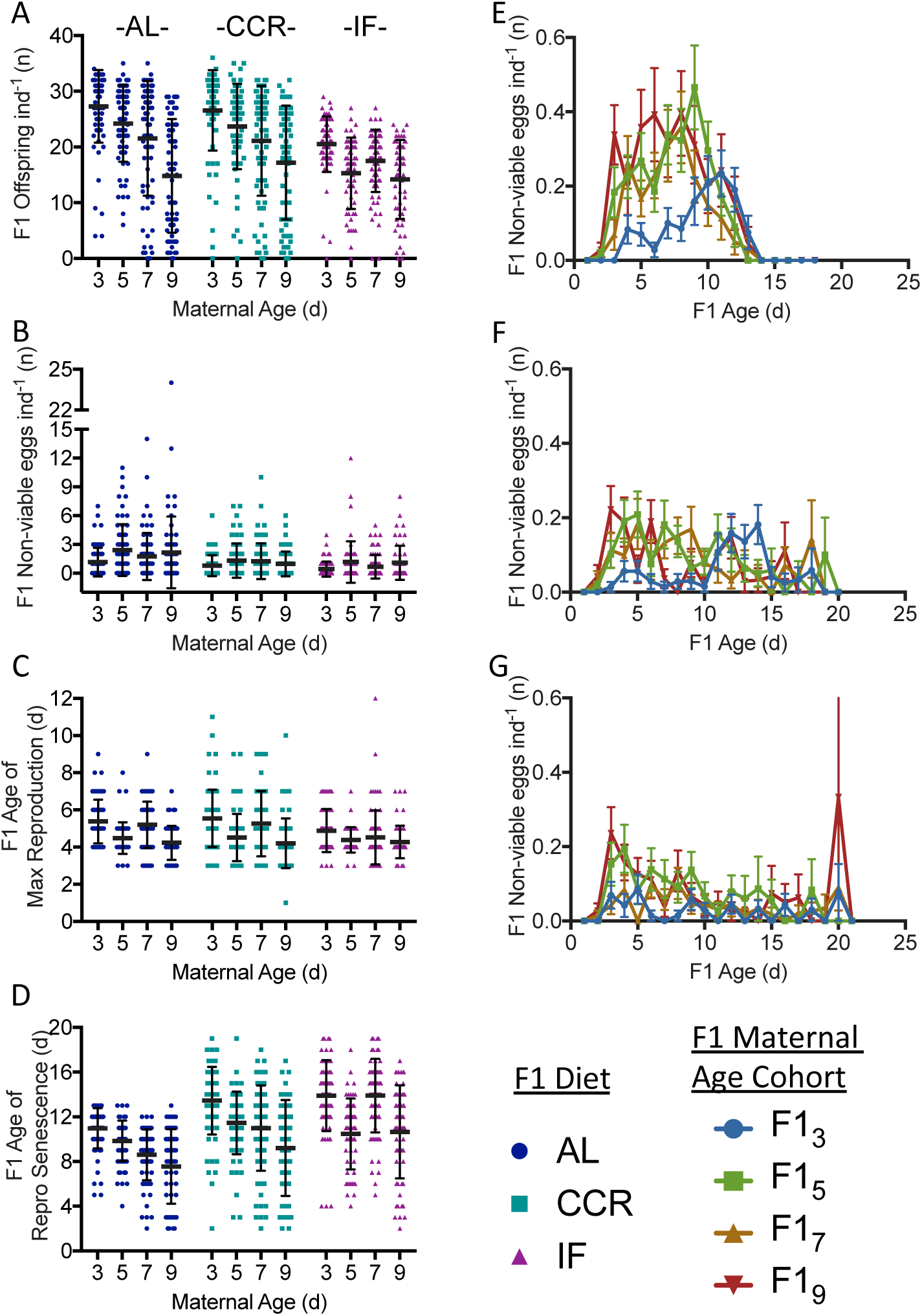
Reproduction in F1s subject to caloric restriction from 3, 5, 7, and 9-d old mothers (F1_3_, F1_5_, F1_7_, and F1_9_, respectively) showing lifetime fecundity **(A)**, number of non-viable offspring **(B)**, age of maximum reproduction (**C**), and reproductive senescence (**D**). Schedule of non-viable offspring production for F1s under *ad libitum* (AL; **E**), chronic caloric restriction (CCR; **F**) and intermittent fasting (IF; **G**) diets. Significance of differences in reproduction is given in Supplementary Table 2. For each F1 maternal age X diet cohort, n = 69 – 72.

Total lifetime fecundity declined significantly with increasing maternal age under all F1 diets (Fig. 4). While lifetime fecundities were similar under AL and CCR for young and middle maternal ages (F1_3_ – F1_7_), for F1_9_ the decline in fecundity under AL was partially rescued by CCR. Maternal age and F1 diet interacted significantly to determine lifetime F1 fecundity (2.0% of the total variance, F_6, 787_ = 3.39, p = 0.0026). Maternal age had a greater impact on fecundity (13.7% of the total variance, F_3, 787_ = 46.62, p < 0.0001) than did F1 diet (7.1% of the variance, F_2, 787_ = 36.04, p < 0.0001).

Non-viable embryo production was strongly influenced by both maternal age (F_3, 785_ = 6.33, p = 0.0003) and F1 diet (F_2, 785_ = 19.32, p < 0.0001) (Fig. 4). The total number of non-viable offspring (unhatched eggs) produced by F1s under AL increased significantly with maternal age, doubling for F1_5_ and F1_9_ (Supplementary Table 2). The timing of non-viable embryo production was strikingly different between the offspring of young and old mothers under AL. For F1_3_, non-viable embryos were low early in life and reached a maximum of 0.2 ind^-1^ d^-1^ late in life at age 11 d. In contrast, non-viable embryos peaked near 0.4 ind^-1^ d^-1^ for F1_5_ and F1_7_ at ages 9 d and 8 d, respectively. For F1_9_, non-viable embryos were produced at a relatively high rate throughout life, peaking at 0.4 ind^-1^d^-1^ at age 6 d. Non-viable embryo production declined significantly under CCR and IF relative to under AL for F1s from older mothers (p < 0.001, except for F1_7_ under CCR, which was not significantly lower) but was still low in early life for offspring of young mothers and high in early life for F1_9_.

Under all food conditions, increasing maternal age significantly decreased the reproductive period and increased the post-reproductive period as a percentage of total lifespan (p < 0.0007 for F1_7_ and F1_9_ relative to F1_3_; Fig. 5, 6; Supplementary Table 3). This effect was greatest for AL and CCR diets, under which the reproductive period was significantly shorter and the post-reproductive period was significantly longer for F1_5_, F1_7_, and F1_9_ than for F1_3_, both in actual days and as a percent of lifespan (Supplementary Table 3, Fig. 5, 6). The pre-reproductive period was not significantly changed by either maternal age or diet except for a slight increase as a percent of total lifespan (though not in actual days) for F1_9_ under AL conditions.

**Fig 5.**
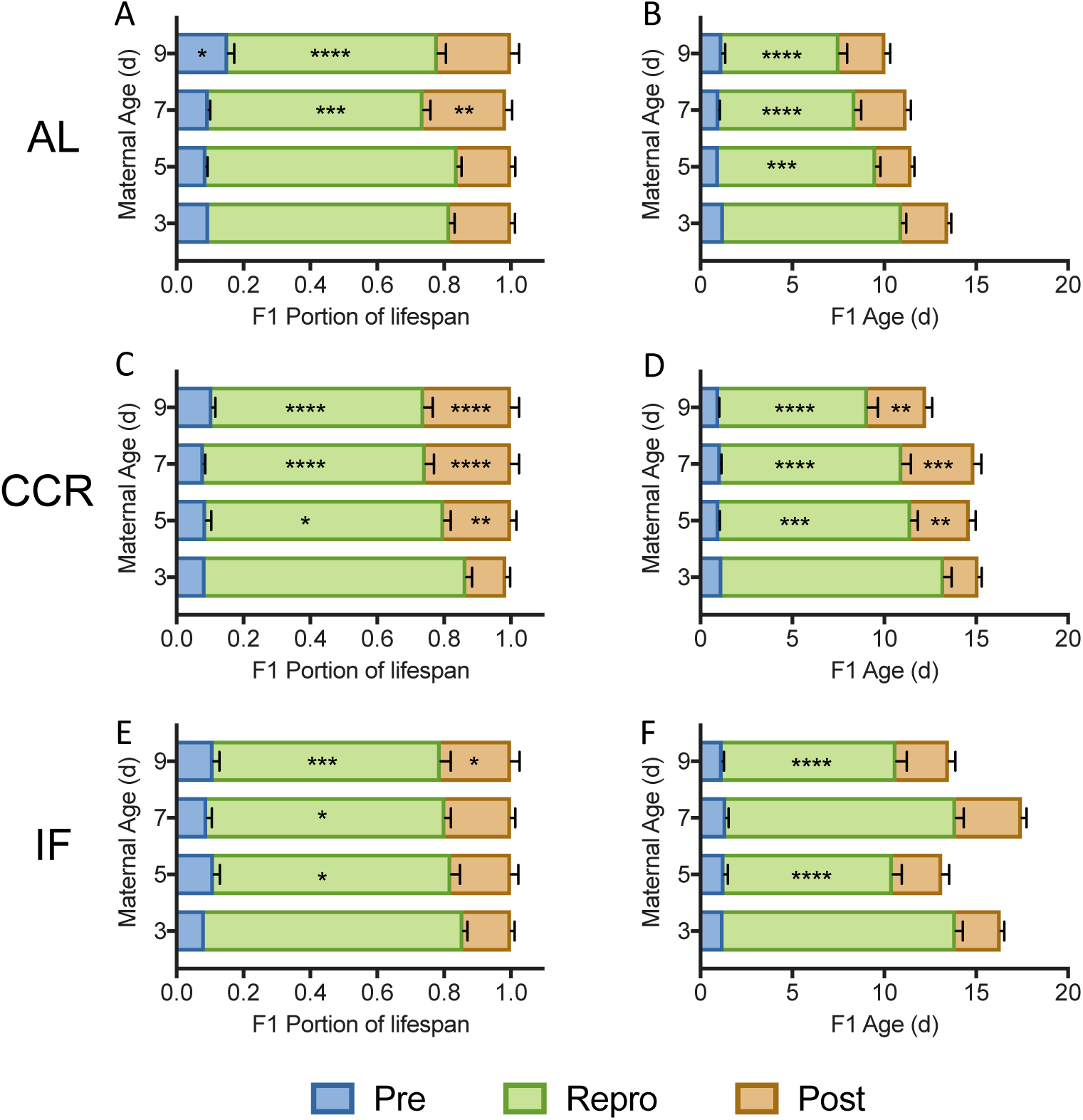
The reproductive schedule of F1s under *ad libitum* (AL; **A-B**), chronic caloric restriction (CCR; **C-D**), or intermittent fasting (IF; **E-F**), shown as a portion of lifespan (left) and as actual days (right). Significant differences in the length of the pre-reproductive, reproductive, and post-reproductive periods in F1_5_– F1_9_ relative to in F1_3_ (Two-way ANOVA with Dunnett’s multiple comparison test) are noted as * (p < 0.05), ** (p < 0.01), *** (p < 0.001), or **** (p < 0.0001). For each F1 maternal age X diet cohort, n = 69 – 72.

**Fig 6.**
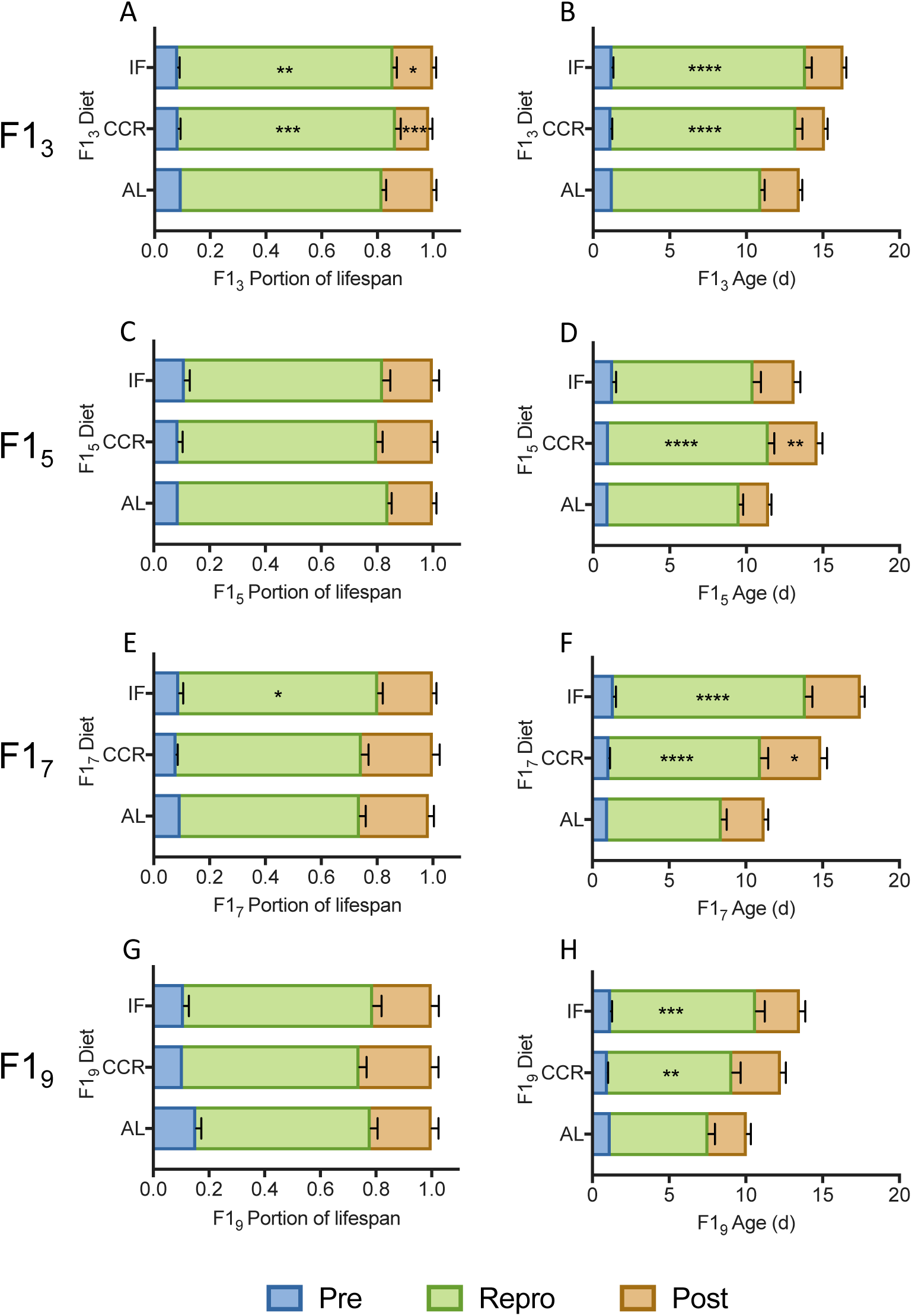
The reproductive schedule of F1s from 3 d (**A-B**), 5 d (**C-D**), 7 d (**E-F**), and 9 d (**G-H**) old mothers, under *ad libitum* (AL), chronic caloric restriction (CCR), or intermittent fasting (IF) diets, shown as a portion of lifespan (left) or as actual days (right). Significant differences in the length of the pre-reproductive, reproductive, and post-reproductive periods in F1_5_ – F1_9_ relative to in F1_3_ (Two-way ANOVA with Dunnetts multiple comparison test) are noted as * (p < 0.05), ** (p < 0.01), *** (p < 0.001), or **** (p < 0.0001). For each F1 maternal age X diet cohort, n = 69 – 72.

Both diet and maternal age changed reproductive continuity (Table 4). Only 2.8% of the F1_3_ AL cohort had discontinuous reproduction; this increased with maternal age to 7.7% for F1_9_ AL, although the difference was not significant. Diet had the greatest effect, with reproductive discontinuity ranging from 12.3 – 17.8% under CCR and 31.5 – 57.7% under IF.

The age of maximum reproduction was generally younger under IF than AL, and for any given diet, the age of maximum reproduction was lower with increasing maternal age (Fig. 4). The interaction between maternal age and diet was not significant (p = 0.08), though both variables significantly impacted age of maximum reproduction independently, with maternal age accounting for 9.7% of the variation (F_3, 777_ = 28.71, p < 0.0001) and diet for 1.4% of the variation (F_2, 777_ = 6.25, p = 0.002). Under AL conditions, the age of maximum reproduction was a full 1.15 days earlier for F1_9_ than for F1_3_. This demonstrates earlier reproductive senescence rather than earlier development, given that there was no difference in the length of time to first reproduction among any maternal age cohorts (Fig. 5). At the peak of reproduction for F1_9_ the number of offspring per individual was already higher for F1_3_; F1_3_ reproduction peaked a day later, when F1_9_ reproduction was already declining.

Maternal age and F1 diet interacted to determine the age of reproductive senescence (Fig. 4), accounting for 3.8% of the total variance (F_6, 778_ = 6.95, p < 0.0001). Diet alone accounted for 11.1% of variance (F_2, 778_ = 60.69, p < 0.0001), and maternal age for 12.3% of the variance (F_3, 778_= 44.71, p < 0.0001). For any given diet, the age of reproductive senescence in F1s was significantly younger with increasing maternal age, except for F1_7_ under IF, in which reproductive senescence was later than for F1_5_ (Fig. 4, Supplementary Table 4). Both CCR and IF significantly delayed reproductive senescence, relative to AL-fed F1s.

### Maternal Investment

We measured female and male egg size and shape, female neonate lipid content, female offspring time to reproductive maturity, and female offspring starvation resistance as estimates of maternal investment in reproduction (Supplementary Fig. 2). As maternal age increased from 3 d to 9 d, female egg area and roundness increased significantly (One-way ANOVA with Tukey’s multiple comparison test, p < 0.0001 for all comparisons). At a maternal age of 11 d, female egg area and roundness significantly decreased. Male eggs showed a similar pattern of increase in size with increasing maternal age and decreased roundness at the oldest maternal age (Supplementary Fig. 2). Neonate lipid content decreased slightly with increasing maternal age, and was only significantly different between F1_3_ and F1_11_ (One-way ANOVA, p = 0.044). We found no significant difference in the time to first reproduction between F1 maternal age or diet cohorts (Fig. 5, 6). Mean lifespan of offspring under starvation conditions decreased significantly with maternal age between F1_3_ and F1_9_, then remained constant for F1_11_ (Supplementary Fig. 2).

### Relative offspring fitness

CCR and IF had different impacts on the correlation between total lifetime fecundity and lifespan. For all F1 maternal age cohorts, the lifespan-reproduction relationship was significantly different between AL and IF (p < 0.0001, extra-sum-of-squares F-test for difference in best-fit values between quadratic equations; Fig. 7). The lifespan-reproduction correlation under CCR was significantly different from under AL for F1_3_ and F1_7_ (p < 0.01) but not for F1_5_ (p = 0.06) or F1_9_ (p = 0.22). The slope for lifespan versus reproduction was much lower under IF than under either AL or CCR, suggesting a greater decrease in lifetime reproduction with increasing lifespan under IF. Under food limitation, the slope of the lifespan-reproduction correlation decreased significantly in the F1_5_, F1_7_, and F1_9_ cohorts relative to the F1_3_ (p < 0.05; extra-sum-of-squares F-test for difference in best-fit values between quadratic equations) suggesting a reduction in fecundity with increasing lifespan under CCR and IF in offspring from older mothers (Fig. 8).

**Fig 7.**
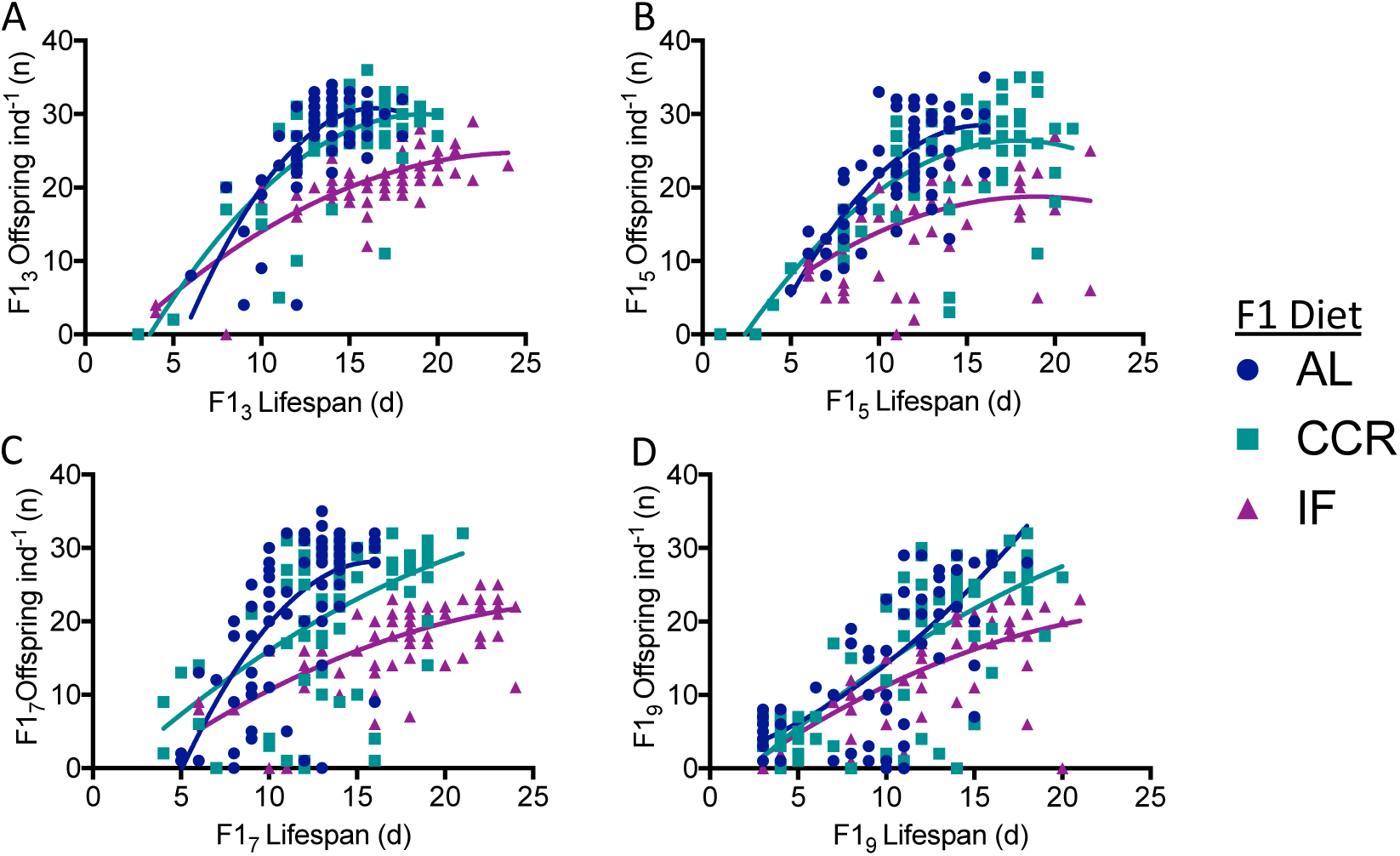
Trade-off between lifespan and lifetime reproduction for F1_3_ (**A**), F1_5_ (**B**), F1_7_ (**C**), and F1_9_ (**D**) under *ad libitum* (AL), chronic caloric restriction (CCR), or intermittent fasting (IF) diets. Relationships are fitted with second-order polynomial (quadratic) equations, and differences between the best-fit values for AL and CCR or AL and IF were determined with an extra-sum-of-squares F-test. CCR was significantly different from AL for F1_3_ and F1_7_ (p < 0.01) but not for F1_5_ (p = 0.06) or F1_9_ (p = 0.22). The regression for IF was significantly different from that for AL for all cohorts (p ≤ 0.0001). For each F1 maternal age X diet cohort, n = 69 – 72.

**Fig 8.**
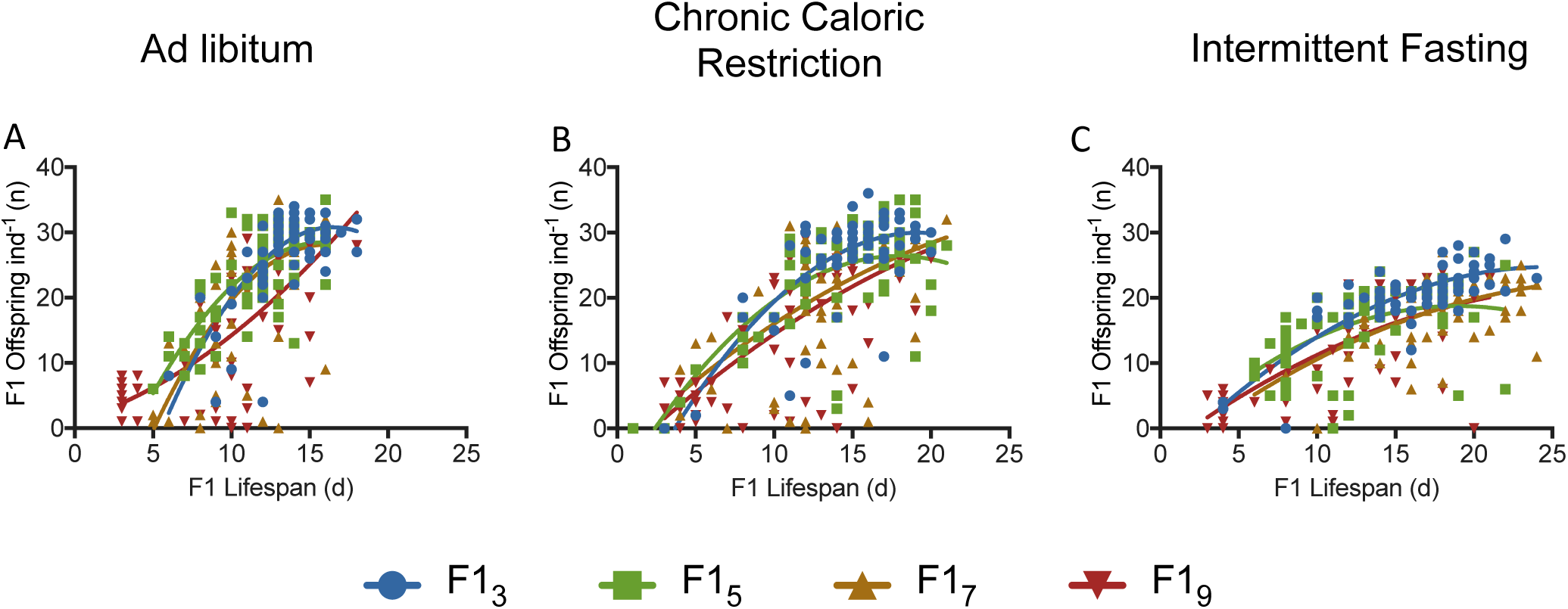
Trade-off between lifespan and lifetime reproduction for F1s from 3, 5, 7, or 9-d old mothers under *ad libitum* (AL; **A**), chronic caloric restriction (CCR; **B**), or intermittent fasting (IF; **C**). Relationships were fitted with second order polynomial (quadratic) equations, and differences between F1_3_ and older maternal age cohorts were tested using an extra-sum-of-squares F-test. Under CCR and IF, the lifespan-reproduction correlation was significantly different for the F1_5_, F1_7_, and F1_9_ cohorts relative to the F1_3_ (p < 0.05). For each F1 maternal age X diet cohort, n = 69 – 72.

We measured age-specific fitness as l_x_m_x_, in which reproduction, l, is multiplied by survivorship, m, for a given day, x. Relative age-specific fitness, defined here as the difference in l_x_m_x_ between calorically restricted and AL-fed rotifers within a maternal age cohort (Fig. 9) or between older maternal age cohorts and F1_3_ for a given diet (Fig. 10), declined with maternal age. Under both CCR and IF, fitness of all F1 maternal age cohorts was much lower in early life relative to AL (Fig. 9). Relative fitness was greater under CCR or IF only late in life, at ages beyond which most AL rotifers were post-reproductive and survivorship was low; this late-life fitness benefit decreased with increasing maternal age. The cumulative relative fitness, measured as net area under the curve, was negative for all maternal age comparisons for a given diet, and for all diet comparisons for a given age, except for F1_9_ CCR relative to F1_9_ AL, which was slightly positive (Fig. 9). The relative fitness under IF was generally lower than that under CCR throughout life. For a given diet, the fitness of older maternal age cohorts relative to F1_3_ was lower throughout life and decreased with increasing maternal age (Fig. 10).

**Fig 9.**
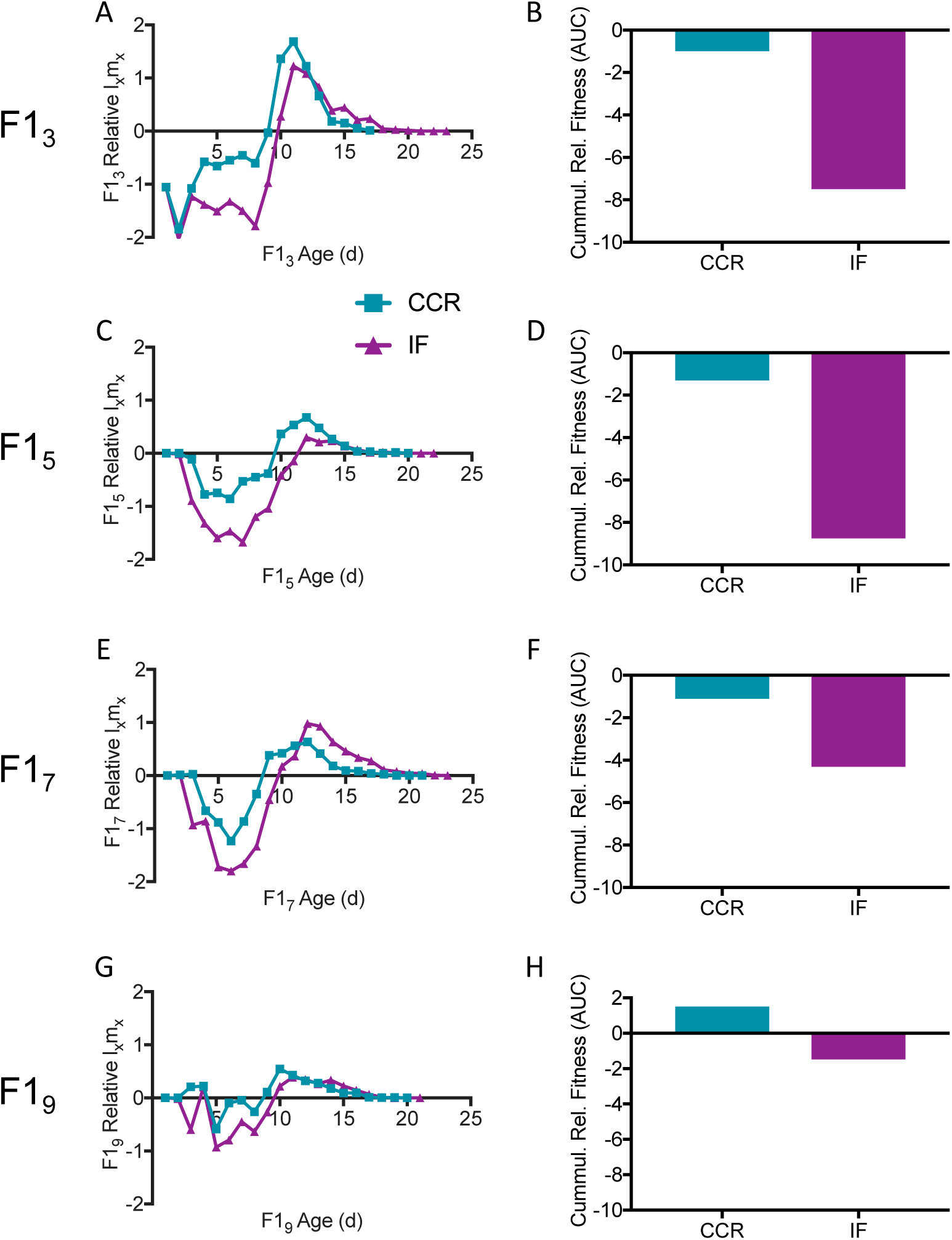
Relative age-specific l_x_m_x_, a measure of relative fitness. Age-specific daily reproduction (l_x_) multiplied by survivorship (m_x_) for F1s under chronic caloric restriction (CCR) or intermittent fasting (IF) is given relative to l_x_m_x_ for *ad libitum* (AL) fed F1s. Relative age-specific fitness (left) and relative cumulative fitness (right) are shown for offspring from different age mothers: (**A-B**) F1_3_, (**C-D**) F1_5_, (**E-F**) F1_7_, (**G-H**) F1_9_. For each F1 maternal age X diet cohort, n = 69 – 72.

**Fig 10.**
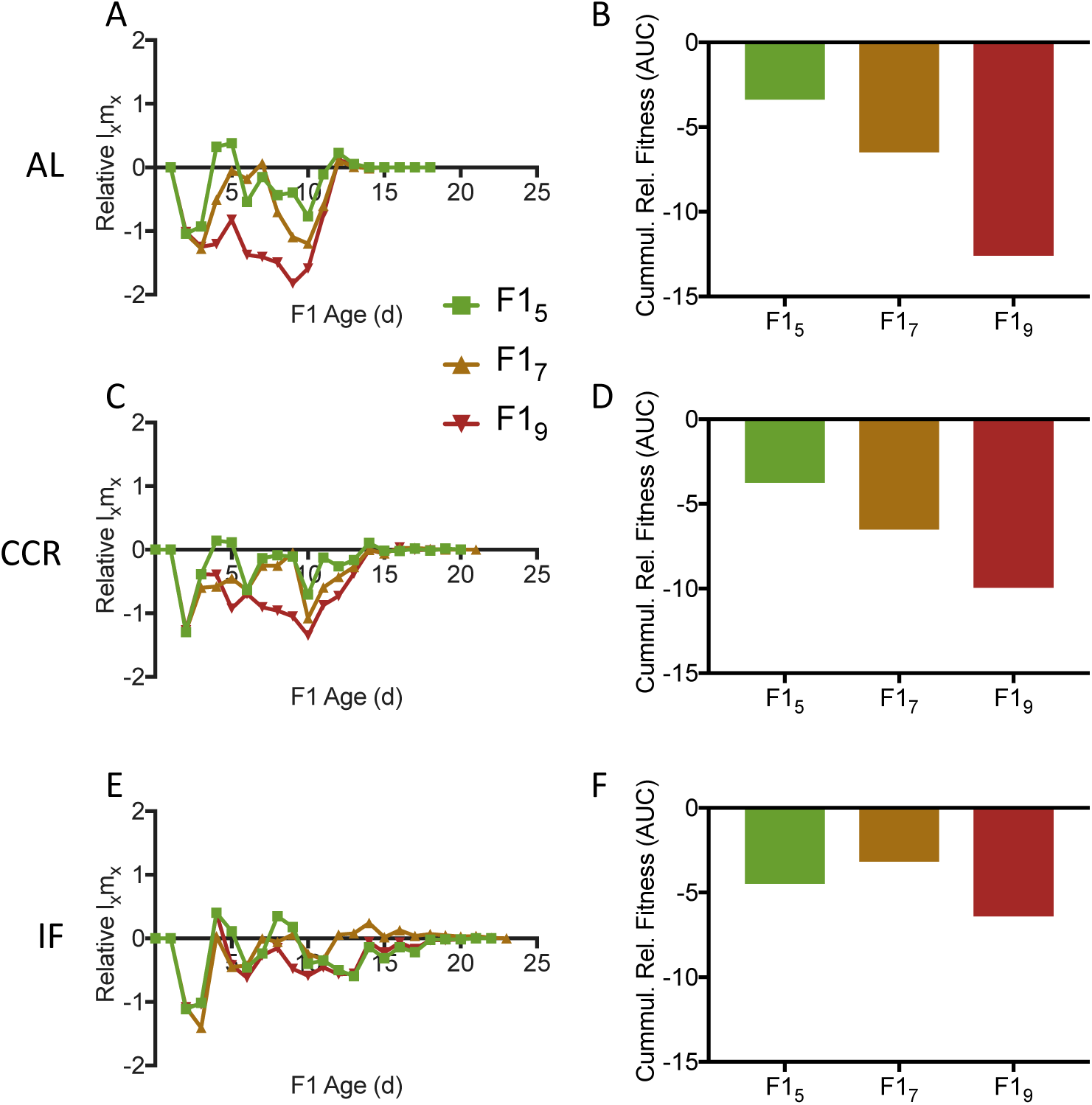
Relative age-specific l_x_m_x_, a measure of relative fitness. Age-specific daily reproduction (l_x_) multiplied by survivorship (m_x_) for offspring of older mothers is given relative to l_x_m_x_ for offspring of the youngest mothers (F1_3_) under *ad libitum* (AL; **A**), chronic caloric restriction (CCR; **C**) and intermittent fasting (IF; **E**). Cumulative relative age-specific fitness (net area under the curve) is shown for AL (**B)**, CCR (**D**), and IF (**F**). For each F1 maternal age X diet cohort, n = 69 – 72.

## DISCUSSION

To our knowledge, this study provides the first evidence that maternal age affects offspring response to caloric restriction, a lifespan extending intervention conserved across a range of taxa. Because maternal environment, physiology, and age are all known to influence offspring phenotype and lifespan, we hypothesized that maternal age might affect offspring adaptive response to caloric restriction. Previous studies have investigated the effect of maternal age on offspring phenotype, but prior work has not examined the combinatorial effect of maternal age, maternal investment, and offspring environment on offspring lifespan, daily and total reproduction, and reproductive schedule. Increasing maternal age changed not only offspring lifespan and the degree of lifespan extension under caloric restriction, but also the length of the reproductive period, fecundity, and the trade-off between lifespan extension and reproduction. The finding that maternal age impacts the magnitude of lifespan extension and the level of resource allocation trade-off has implications for the widespread use of CR and CR mimetics as anti-aging therapies in humans, and should be verified in mammalian models of aging.

### Increasing maternal age decreases offspring lifespan and fecundity

Consistent with earlier studies in rotifers and other species, offspring of the oldest mothers had a significantly shorter median lifespan than the offspring from the youngest mothers 23,28,29,31,33,40,81,82. Among AL-fed F1s, earlier onset of aging, rather than an increased rate of aging, appeared to be responsible for the observed decrease in lifespan in old-mother offspring.

The decline in offspring fecundity with increasing maternal age found in this and previous rotifer studies ^23,76^ differs from some reports for *C. elegans, Daphnia*, and *Drosophila,* in which offspring from the youngest or smallest mothers have been shown to have lower lifetime reproduction than those from older mothers ^34,40,83^. One possible explanation is that differences in reproductive strategy may drive the differences in offspring outcomes with changing maternal age among varied small, short-lived invertebrate species. *Brachionus manjavacas* makes a relatively large investment in each offspring, producing a maximum of 25 – 30 eggs over its lifespan, with each embryo approximately one-third the size of its mother. In comparison, hermaphroditic *C. elegans* produces up to 300 offspring of only 30 – 50 μm in size over a shorter reproductive period, laying up to 140 eggs per day ^72,74^. *Drosophila* lay up to 100 eggs per day with approximately 600 total offspring, and *Daphnia* produce nearly 100 offspring in multiple synchronized batches that are coordinated with adult molting ^73,84,85^. Additionally, these other invertebrates are indirect developers, producing offspring with larval stages or that undergo multiple metamorphoses before becoming reproductively mature; *B. manjavacas,* in contrast, has direct development, with neonates emerging from the egg as a small version of the adult form.

The early, high-investment, and direct reproductive strategy may be adaptive for *B. manjavacas*, although it is different from the r-selection strategy expected for a microscopic invertebrate that evolved in ephemeral habitats where it was subject to high extrinsic mortality due to predation and rapidly changing environmental conditions ^86^. A larger investment in each embryo increases chances of neonate survival, but high external mortality likely decreases the selection pressure to produce high-quality offspring at late maternal ages ^63^. Differences in life history strategy, even among short-lived invertebrate models evolving under similar environmental and predation selective pressures, must be considered when determining the applicability of results among species and from model organisms to humans.

### Offspring fitness is not determined by simple changes in gross maternal resource allocation

This study suggests that offspring size is not a sufficient measure to determine the quality or quantity of maternal investment in reproduction. Despite larger egg size and neonate body size 76, we did not observe the accelerated development time or greater early-life reproductive output that has been associated with earlier onset of senescence in *Daphnia* old-mother offspring ^40^. Lipid reserves and starvation resistance were slightly lower in old-mother offspring, suggesting decreased offspring provisioning, though likely not enough to account for the 21% reduction in lifespan and 46% decline in reproduction between the youngest mother and oldest mother offspring. The relatively synchronous time to death in maternal age cohorts under starvation conditions suggests that maternal provisioning to offspring is relatively consistent among offspring for a given maternal age cohort, but may decrease slightly with increasing maternal age. Lifespan extension under CCR and IF demonstrates that lifespan is plastic for all maternal age F1 cohorts, and is not solely determined by maternal provisioning. Taken together, these findings suggest that epigenetic or cellular mechanisms beyond simple changes in gross maternal investment play a role in decreased offspring fitness with increasing maternal age.

### Maternal age alters offspring response to caloric restriction

The maternal age of the experimental cohort changes the magnitude of lifespan extension and degree of reproductive trade-off in response to caloric restriction, and may thus change interpretation of the mechanism. Lifespan extension under caloric restriction was greater for the offspring of the oldest mothers; under IF relative to AL, less trade-off between lifespan and reproduction was observed for F1_9_ (no reduction in mean lifetime reproduction for a 28% increase in lifespan) than for F1_3_ (35% reduced net reproduction and 21% lifespan increase). A previous study in *B. manjavacas* similarly showed that old-mother offspring had greater lifespan extension when their mothers were calorically restricted ^76^. While the rate of aging decreased under caloric restriction in young mother offspring, only the onset of aging and not the aging rate were altered in old mother offspring. These results suggest that the offspring of the youngest mothers may already be closer to potential maximum lifespan, or alternatively, are less able to up-regulate caloric restriction-induced protective pathways. Given that F0 survivorship was 67% at 9 d old, we cannot rule out that a change in phenotypic composition of the population due to mortality led to the observed changes in offspring lifespan, fecundity, and caloric restriction response. However, as the tested rotifer population was isogenic and no other external environmental variables changed over the course of the experiment, the observed differences in both magnitude and mechanism of the response to caloric restriction are likely due to maternal age. Increasing maternal age leads to changes in offspring gene expression in *C. elegans*, which likely cause differential offspring responses to environmental conditions ^83^. The maternal age of experimental cohorts is not always controlled or consistent among separate aging studies and thus maternal age effects may be a source of the observed variability and inconsistencies seen among caloric restriction experiments.

### Caloric restriction increases relative fitness only in late life

To assess the effects of maternal age and to evaluate the Disposable Soma theory and the evolutionary theory of the response to caloric restriction, many studies focus on end-point assessments such as median and maximum lifespan and on trade-offs between longevity and lifetime reproduction ^29,87-90^. In reality, evolutionary fitness is an age-specific combination of innate lifespan, resistance to external mortality, and reproduction. Relative l_x_m_x_ provides an age-specific measure of relative fitness that incorporates age-specific survivorship, fecundity, latency to reproduction, and timing of reproductive senescence.

Averaged over lifetime, the shift to lower daily reproduction and extended lifespan under CCR and IF appears maladaptive relative to the reproductive strategy under AL; the integrated relative lifetime l_x_m_x_ is negative for CCR and IF. CCR and IF both provide an age-specific late life benefit, however, supporting the Disposable Soma theory for the evolution of lifespan extension in response to caloric restriction. It is hypothesized that those individuals that are able to reallocate resources from reproduction to maintenance of the soma during times of famine have a selective advantage; this strategy allows the organism to make it through the period of starvation, and produce offspring later when resources become available ^55,59^. In the current study, the relationship between lifespan and lifetime reproduction was positive within each population, showing that longer-lived individuals tended to reproduce more. When food was limited by CCR or IF, however, the slope of the lifespan-reproduction correlation decreased, suggesting a trade-off between extending lifespan and producing offspring when resources are limited, as is the expectation under the Disposable Soma theory ^55^.

The caloric restriction-mediated late-life relative l_x_m_x_ benefit greatly declines for F1s with increasing maternal age. Given that offspring of older mothers had a proportionally greater increase in lifespan, this does not indicate a decreased selective pressure for an adaptive response to caloric restriction. Rather, it suggests that the overall decreased lifespan and fecundity in old-mother offspring negates any fitness benefit of the caloric restriction response. While the early life cost of caloric restriction appears to decline with increasing maternal age, this is due primarily to the decrease in l_x_m_x_ under AL conditions throughout life with increasing maternal age. The decline in relative offspring fitness with increasing maternal age supports the hypothesis that the force of natural selection decreases with increasing age ^63,91,92^.

The increase in lifespan with concomitant decrease in daily reproduction is not in itself an adaptive response that increases fitness. Indeed, fitness will only be increased if reproduction is upregulated once food is restored. The ability to re-establish reproduction in late life after early life caloric restriction should be tested in the context of maternal age. Given the low rates of reproduction in old-mother offspring even under full food conditions, it is unclear that there is an evolutionary benefit to increasing lifespan in the face of limiting food resources for old-mother offspring, rather than maximizing early reproduction.

## CONCLUSIONS

Because the selection gradient on both mortality and fecundity are decreasing with increasing age, changes in these parameters that affect older age classes are theorized to have less impact than the same changes at earlier ages ^63,92-94^. This study provides empirical support for this hypothesis, and demonstrates that maternal age affects not only offspring fitness, but also offspring response to interventions. We observed changing levels of caloric restriction-mediated lifespan extension and reproductive trade-off in different maternal age offspring. Additional work is needed to determine if maternal age has a similar impact on other lifestyle, diet, or pharmaceutical interventions, or if there are differences in maternal age effects among varied genotypes. Controlling for maternal age in experimental populations will be important for replication of experimental results, appropriate interpretation of findings, and assignment of mechanism.

## Authors’ contributions

K.E.G. designed and supervised the experiments, interpreted the data, and wrote the manuscript. M.J.B., G.J., E.C., and E.S conducted the experiments and edited the manuscript.

## Data availablility

Upon publication, data will be included in online supplementary material.

## Acknowledgements

We thank Michael Neubert, Hal Caswell, Christina Hernandez, and Silke vanDaalen for discussions and constructive comments on the manuscript. We appreciate suggestions from anonymous reviewers that improved the manuscript.

