## Supplementary Information for "Maternal age alters offspring lifespan, fitness, and lifespan extension under caloric restriction"

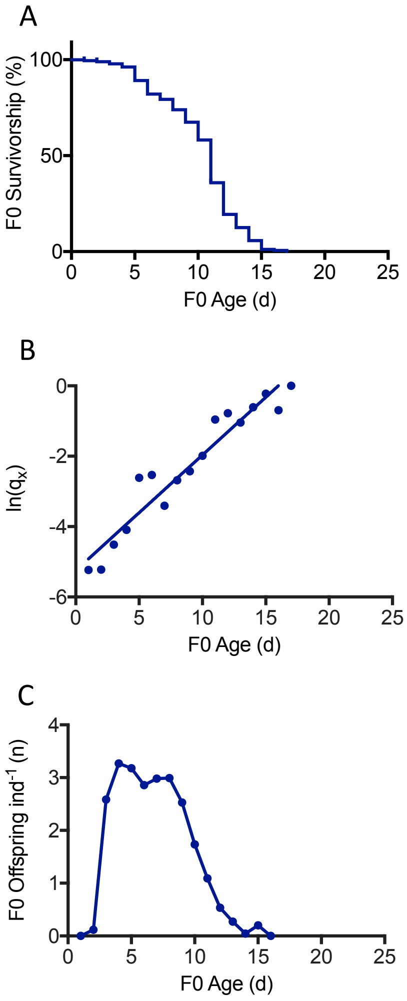


Supplementary Fig. 1. Maternal (F0) survivorship **(A)**, hazard rate **(B)**, and daily reproduction rate **(C)**. n = 187.

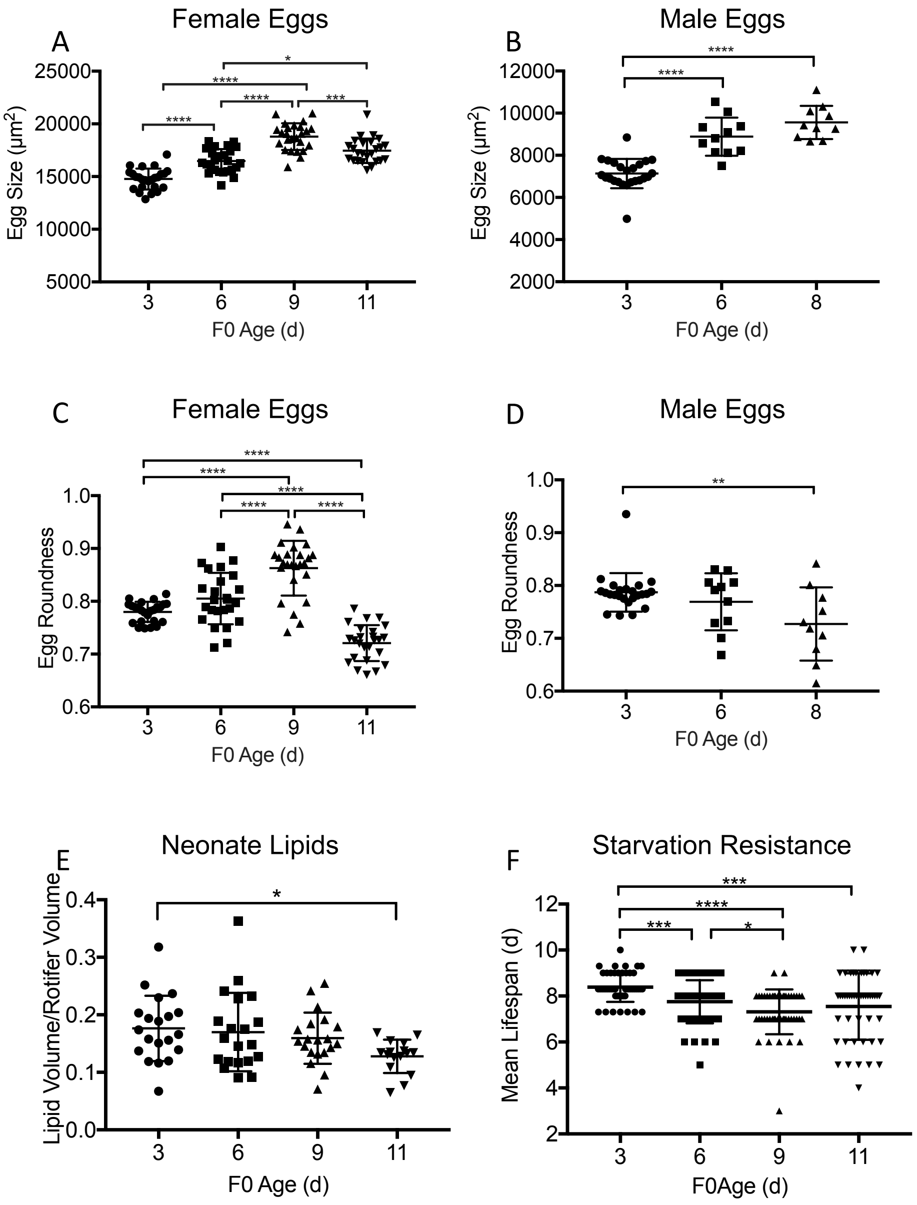


Supplementary Fig. 2. Maternal investment in offspring. Amictic egg (hatch into females, n = 20) and mictic egg (hatch into males, n = 10) size (**A**, and **B**, respectively) and shape (**C** and **D**) from F0 mothers of different ages. All multiple comparisons between every maternal age were significantly different for egg size and shape except for maternal age 3 d versus 6 d for both amictic and mictic egg roundness (one-way ANOVA with Tukey's test for multiple comparisons). Neonate lipid content **(E)** was significantly different only between maternal ages of 3 d and 11 d ($p = 0.04$; one-way ANOVA with Holm-Sidak's test for multiple comparisons, n = 20). Offspring starvation resistance **(F)** was significantly different among all maternal ages except 6 d and 9 d versus 11 d (n = 48). Significance indicated as * ($p < 0.05$), ** ($p < 0.01$), *** ($p < 0.001$), or **** ($p < 0.0001$).

| | | LIFESPAN | | | | | GOMPERTZ | | | | | |
|---|---|---|---|---|---|---|---|---|---|---|---|---|
| Maternal Age Cohort | Diet | Median Lifespan (d) | Percent Change from AL | Median Diff than AL? (p) | Median Diff than $F1_3$? (p) | Maximum lifespan (d; 95%) | α (intercept) | α Diff than AL? (p) | β (slope) | β Diff than AL? (p) | $R^2$ | n |
| **F0** | **AL** | 11 | | | | | -5.247 ± 0.2523 | | 0.3281 ± 0.02462 | | 0.922 | 187 |
| | **AL** | 14 | | | | 16.4 | -6.995 ± 0.5566 | | 0.3959 ± 0.04306 | | 0.894 | 71 |
| **$F1_3$** | **CCR** | 16 | 14.3 | **<0.0001** | | 19.35 | -5.714 ± 0.3134 | ‡ | 0.2677 ± 0.04306 | **0.0109** | 0.921 | 72 |
| | **IF** | 17 | 21.4 | **<0.0001** | | 21.35 | -5.478 ± 0.488 | ‡ | 0.2251 ± 0.04306 | **0.0047** | 0.815 | 72 |
| | **AL** | 12 | | | **<0.0001** | 16 | -6.169 ± 0.7776 | | 0.3715 ± 0.06794 | | 0.769 | 72 |
| **$F1_5$** | **CCR** | 15 | **25.0** | **<0.0001** | 0.9490 | 20 | -5.727 ± 0.3153 | **0.0038** | 0.266 ± 0.0232 | 0.101 | 0.910 | 71 |
| | **IF** | 13 | **8.3** | **0.0004** | **0.0011** | 20 | -4.508 ± 0.5135 | ‡ | 0.1855 ± 0.03497 | **0.0188** | 0.684 | 71 |
| | **AL** | 12 | | | **<0.0001** | 16 | -5.395 ± 0.5705 | | 0.3151 ± 0.05162 | | 0.788 | 71 |
| **$F1_7$** | **CCR** | 14 | 16.7 | **<0.0001** | 0.1999 | 21 | -5.422 ± 0.3754 | **0.0056** | 0.2545 ± 0.02842 | 0.3001 | 0.851 | 69 |
| | **IF** | 18 | **50.0** | **<0.0001** | **0.0183** | 24 | -5.784 ± 0.4458 | **<0.0001** | 0.2192 ± 0.02771 | 0.1021 | 0.817 | 72 |
| | **AL** | 11 | | | **<0.0001** | 16 | -4.179 ± 0.4649 | | 0.229 ± 0.04152 | | 0.717 | 72 |
| **$F1_9$** | **CCR** | 13 | 18.2 | **0.0003** | **0.0151** | 19.7 | -4.037 ± 0.3194 | **0.016** | 0.1653 ± 0.02428 | 0.1776 | 0.768 | 72 |
| | **IF** | 14 | 27.3 | **<0.0001** | **0.0001** | 20 | -4.804 ± 0.4156 | **0.0016** | 0.211 ± 0.02946 | 0.7222 | 0.786 | 72 |

Supplementary Table 1. Changes in lifespan and mortality rate in maternal females (F0) and in offspring from 3, 5, 7, and 9-d old mothers ($F1_3$, $F1_5$, $F1_7$, and $F1_9$, respectively) under *ad libitum* (AL; 6 x $10^5$ cells $ml^{-1}$ *Tetraselmis suecica*), chronic caloric restriction (CCR; 6 x $10^4$ cells $ml^{-1}$ *T. suecica*, a 90% reduction in food relative to AL), or intermittent fasting (IF; alternate day AL and starvation) diets. Significant differences are shown in bold. ‡ Because slopes differ so much, it is not possible to test whether the intercepts differ significantly. Column 4 shows where the percent change from AL is significantly different from that for $F1_3$ in bold (z test for two population proportions, $p < 0.05$).

| | Diet | Net Repro Rate (Ro) | Diff than AL? (p) | Diff than $F1_3$? (p) | Max Daily Repro | Diff than AL? (p) | Diff than $F1_3$? (p) | Non-viable eggs (n) | Diff than AL? (p) | Diff than $F1_3$? (p) |
|---|---|---|---|---|---|---|---|---|---|---|
| | **AL** | 27.30 ± 0.77 | | | 3.60 ± 0.12 | | | 1.15 ± 0.18 | | |
| **$F1_3$** | **CCR** | 26.56 ± 0.85 | **<0.0001** | | 3.14 ± 0.09 | **0.0010** | | 0.79 ± 0.13 | 0.5221 | |
| | **IF** | 20.5 ± 0.59 | **<0.0001** | | 2.31 ± 0.07 | **<0.0001** | | 0.43 ± 0.09 | 0.0773 | |
| | **AL** | 24.22 ± 0.86 | | 0.1145 | 3.73 ± 0.11 | | 0.8088 | 2.42 ± 0.34 | | **0.0014** |
| **$F1_5$** | **CCR** | 23.68 ± 0.97 | **<0.0001** | 0.1586 | 3.13 ± 0.10 | **<0.0001** | 0.9609 | 1.30 ± 0.22 | **0.0048** | 0.4502 |
| | **IF** | 15.29 ± 0.86 | **<0.0001** | **0.0016** | 2.38 ± 0.10 | **<0.0001** | 0.6403 | 1.16 ± 0.29 | **0.0019** | 0.1709 |
| | **AL** | 21.56 ± 1.22 | | **0.0001** | 3.49 ± 0.19 | | 0.8088 | 1.75 ± 0.29 | | 0.2913 |
| **$F1_7$** | **CCR** | 21.11 ± 1.178 | **0.0074** | **0.0003** | 2.75 ± 0.16 | **0.0018** | **0.0017** | 1.25 ± 0.22 | 0.3057 | 0.5106 |
| | **IF** | 17.52 ± 0.66 | **0.0205** | 0.1159 | 2.13 ± 0.11 | **<0.0001** | 0.5060 | 0.68 ± 0.15 | **0.0042** | 0.883 |
| | **AL** | 14.82 ± 1.25 | | **<0.0001** | 3.06 ± 0.22 | | 0.0618 | 2.17 ± 0.47 | | **0.017** |
| **$F1_9$** | **CCR** | 17.20 ± 1.26 | 0.1998 | **<0.0001** | 2.53 ± 0.14 | 0.0529 | **0.0011** | 0.97 ± 0.16 | **0.0018** | 0.9535 |
| | **IF** | 14.20 ± 0.95 | 0.9023 | **<0.0001** | 2.49 ± 0.13 | 0.0529 | 0.5060 | 1.09 ± 0.24 | **0.0088** | 0.2503 |

Supplementary Table 2. Changes in reproduction in offspring from 3, 5, 7, and 9-d old mothers ($F1_3$, $F1_5$, $F1_7$, and $F1_9$, respectively) under *ad libitum* (AL; 6 x $10^5$ cells $ml^{-1}$ *Tetraselmis suecica*), chronic caloric restriction (CCR; 6 x $10^4$ cells $ml^{-1}$ *T. suecica*, a 90% reduction in food relative to AL), or intermittent fasting (IF; alternate day AL and starvation) diets. Significant differences ($p < 0.05$) are shown in bold.

| | Diet | Pre-Repro Period (d) | Diff than AL? (p) | Diff than $F1_3$? (p) | Repro Period (d) | Diff than AL? (p) | Diff than $F1_3$? (p) | Post-Repro Period (d) | Diff than AL? (p) | Diff than $F1_3$? (p) |
|---|---|---|---|---|---|---|---|---|---|---|
| | **AL** | 1.28 ± 0.05 | | | 9.69 ± 0.22 | | | 2.51 ± 0.17 | | |
| **$F1_3$** | **CCR** | 1.19 ± 0.05 | 0.7016 | | 12.07 ± 0.40 | **0.0003** | | 1.86 ± 0.18 | **0.0001** | |
| | **IF** | 1.26 ± 0.06 | 0.6308 | | 12.66 ± 0.38 | **0.0019** | | 2.44 ± 0.19 | **0.0231** | |
| | **AL** | 1.00 ± 0.0 | | 0.9680 | 8.56 ± 0.23 | | 0.3610 | 1.92 ± 0.16 | | 0.6006 |
| **$F1_5$** | **CCR** | 1.03 ± 0.03 | 0.9993 | 0.9995 | 10.43 ± 0.37 | 0.1913 | **0.0199** | 3.19 ± 0.32 | 0.1787 | **0.0049** |
| | **IF** | 1.30 ± 0.19 | 0.6206 | 0.5925 | 9.17 ± 0.48 | 0.1946 | **0.0347** | 2.68 ± 0.38 | 0.7035 | 0.3462 |
| | **AL** | 1.03 ± 0.03 | | 0.9999 | 7.39 ± 0.32 | | **0.0007** | 2.79 ± 0.24 | | **0.0062** |
| **$F1_7$** | **CCR** | 1.11 ± 0.04 | 0.8261 | 0.9959 | 9.87 ± 0.46 | 0.6690 | **<0.0001** | 3.92 ± 0.38 | 0.9438 | **<0.0001** |
| | **IF** | 1.41 ± 0.13 | 0.9819 | 0.9819 | 12.49 ± 0.43 | **0.0114** | **0.0258** | 3.58 ± 0.26 | 0.0886 | 0.0570 |
| | **AL** | 1.19 ± 0.16 | | 0.0253 | 6.37 ± 0.43 | | 0.6006 | 2.51 ± 0.27 | | 0.2180 |
| **$F1_9$** | **CCR** | 1.02 ± 0.02 | 0.2879 | 0.7595 | 8.11 ± 0.54 | 0.9771 | **<0.0001** | 3.17 ± 0.32 | 0.3940 | **<0.0001** |
| | **IF** | 1.21 ± 0.07 | 0.3804 | 0.5925 | 9.46 ± 0.57 | 0.2504 | **0.0005** | 2.86 ± 0.38 | 0.9625 | **0.0181** |

Supplementary Table 3. Changes in length of pre-reproductive, reproductive, and post-reproductive periods in offspring from 3, 5, 7, and 9-d old mothers ($F1_3$, $F1_5$, $F1_7$, and $F1_9$, respectively) under *ad libitum* (AL; 6 x $10^5$ cells $ml^{-1}$ *Tetraselmis suecica*), chronic caloric restriction (CCR; 6 x $10^4$ cells $ml^{-1}$ *T. suecica*, a 90% reduction in food relative to AL), or intermittent fasting (IF; alternate day AL and starvation) diets. Significant differences ($p < 0.05$) are shown in bold.

| | Diet | Age at Max. Repro (d) | Diff than AL? (p) | Diff than $F1_3$? (p) | Age at Repro Senesc. (d) | Diff than AL? (p) | Diff than $F1_3$? (p) | Discont. Repro Period (%) | Diff than AL? (z, p) | Diff than $F1_3$? (z, p) |
|---|---|---|---|---|---|---|---|---|---|---|
| | **AL** | 5.38 ± 0.14 | | | 10.97 ± 0.21 | | | 2.8 | | |
| **$F1_3$** | **CCR** | 5.55 ± 0.18 | 0.697 | | 13.45 ± 0.36 | **<0.0001** | | 17.8 | 2.94, **0.0032** | |
| | **IF** | 4.89 ± 0.14 | **0.048** | | 13.89 ± 0.37 | **<0.0001** | | 45.8 | 5.98, **<0.0001** | |
| | **AL** | 4.49 ± 0.11 | | **0.0002** | 9.84 ± 0.23 | | 0.163 | 6.3 | | 0.97, 0.333 |
| **$F1_5$** | **CCR** | 4.52 ± 0.16 | 0.989 | **<0.0001** | 11.46 ± 0.35 | **0.0112** | **0.0016** | 15.9 | 2.99, 0.083 | 0.09, 0.76 |
| | **IF** | 4.39 ± 0.09 | 0.909 | 0.1187 | 10.47 ± 0.44 | 0.5325 | **<0.0001** | 38.9 | 4.32, **<0.0001** | 0.61, 0.780 |
| | **AL** | 5.20 ± 0.15 | | 0.825 | 8.60 ± 0.27 | | **<0.0001** | 4.3 | | 0.47, 0.64 |
| **$F1_7$** | **CCR** | 5.26 ± 0.21 | 0.955 | 0.5173 | 10.99 ± 0.45 | **<0.0001** | **<0.0001** | 14.1 | 4.04, **0.044** | 0.37, 0.54 |
| | **IF** | 4.52 ± 0.17 | **0.004** | 0.3041 | 13.90 ± 0.39 | **<0.0001** | >0.9999 | 57.8 | 6.85, **<0.0001** | 1.43, 0.154 |
| | **AL** | 4.23 ± 0.11 | | **<0.0001** | 7.56 ± 0.42 | | **<0.0001** | 7.8 | | 1.23, 0.199 |
| **$F1_9$** | **CCR** | 4.21 ± 0.16 | 0.994 | **<0.0001** | 9.22 ± 0.53 | **0.0085** | **<0.0001** | 12.3 | 0.88, 0.381 | 0.81, 0.37 |
| | **IF** | 4.28 ± 0.12 | 0.977 | **0.0342** | 10.66 ± 0.56 | **<0.0001** | **<0.0001** | 31.5 | 3.33, **0.0009** | 1.63, 0.103 |

Supplementary Table 4. Changes in ages of maximum reproduction and reproductive senescence and in the continuity of the reproductive period in offspring from 3, 5, 7, and 9-d old mothers ($F1_3$, $F1_5$, $F1_7$, and $F1_9$, respectively) under *ad libitum* (AL; 6 x $10^5$ cells $ml^{-1}$ *Tetraselmis suecica*), chronic caloric restriction (CCR; 6 x $10^4$ cells $ml^{-1}$ *T. suecica*, a 90% reduction in food relative to AL), or intermittent fasting (IF; alternate day AL and starvation) diets. Significant differences ($p < 0.05$) are shown in bold.
